# The functional organization of excitation and inhibition in the dendritic arbors of retinal direction-selective ganglion cells

**DOI:** 10.1101/718783

**Authors:** Varsha Jain, Benjamin L. Murphy-Baum, Geoff deRosenroll, Santhosh Sethuramanujam, Mike Delsey, Kerry Delaney, Gautam B. Awatramani

## Abstract

Recent studies indicate that the precise timing and location of excitation and inhibition (E/I) within active dendritic trees can significantly impact neuronal function. How excitatory and inhibitory inputs are functionally organized at the subcellular level in intact circuits remains unclear. To address this issue, we took advantage of the retinal direction-selective ganglion cell circuit, in which directionally tuned inhibitory GABAergic input arising from starburst amacrine cells shape direction-selective dendritic responses. We combined two-photon Ca^2+^ imaging with genetic, pharmacological, and single-cell ablation methods to examine local E/I. We demonstrate that when active dendritic conductances are blocked, direction selectivity emerges semi-independently within unusually small dendritic segments (<10 µm). Impressively, the direction encoded by each segment is relatively homogenous throughout the ganglion cell’s dendritic tree. Together the results demonstrate a precise subcellular functional organization of excitatory and inhibitory input, which suggests that the parallel processing scheme proposed for direction encoding could be more fine-grained than previously envisioned.

## INTRODUCTION

Neural computations often rely on interactions between excitatory and inhibitory synaptic inputs (Cafaro and Rieke, 2010; D’amour and Froemke, 2015; Denève and Machens, 2016; Rupprecht and Friedrich, 2018; Wehr and Zador, 2003; Xue et al., 2014). While excitation is required for driving action potential firing, inhibition serves as an opposing force to gate excitatory activity (Grienberger et al., 2017; Isaacson and Scanziani, 2011; Koch et al., 1982; Lovett-Barron et al., 2012; Muñoz et al., 2017; Murayama et al., 2009; Poleg-Polsky et al., 2018; Ranganathan et al., 2018). Such interactions between excitation and inhibition (E/I) form the basis for logical ‘AND-NOT’ computations, in which activity is only propagated when excitation is present but inhibition is not (Barlow and Levick, 1965; Major et al., 2008; Schachter et al., 2010; Sivyer and Williams, 2013; Wilson et al., 2018). The direction-selective ganglion cell (DSGC) circuit in the retina is a classic example of a circuit that performs AND-NOT computations: objects in the visual scene that move over a DSGC’s receptive field in its ‘preferred’ direction drive excitation onto its dendrites that trigger action potential firing, but for motion in the opposite or ‘null’ direction, excitation is vetoed by coincident GABAergic inhibition (**Figure 1A-B**).

**Figure 1.**
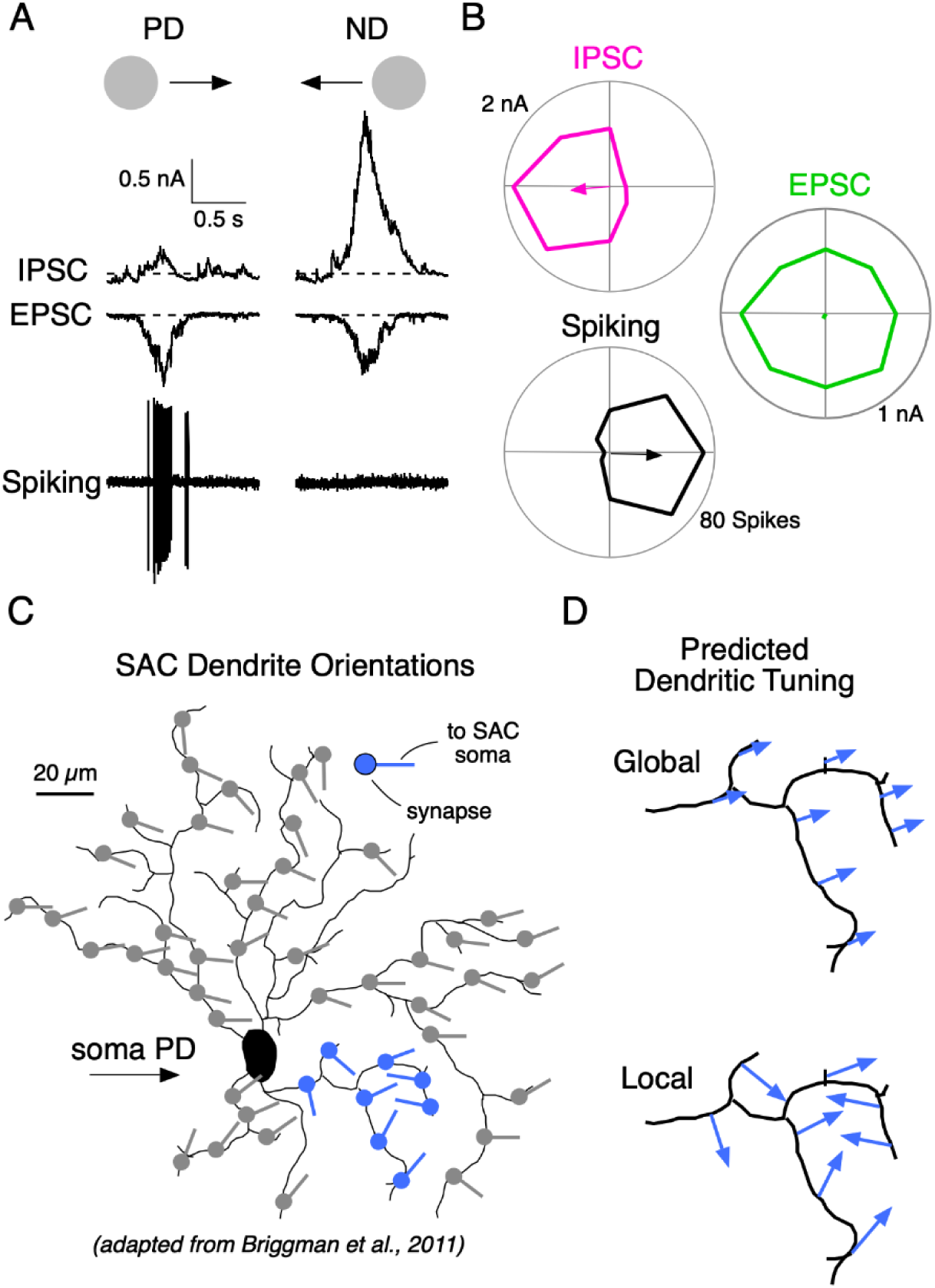
Dendritic direction selectivity is determined by the relative strength of local excitatory and inhibitory inputs and the spatial scale of dendritic integration. **A**, spiking, IPSCs and EPSCs recorded in succession from the soma of a single ON-OFF DSGC in response to a positive contrast spot (200 µm diameter; 500 µm/s) moving in the DSGC’s preferred or null direction (PD and ND). **B**, Polar plots of the peak amplitudes of the IPSC, EPSCs and spiking responses evoked by spots moving in 8 directions. **C**, Schematic of a reconstructed ON-OFF DSGC showing that the average orientation of starburst amacrine cell (SAC) dendrites is predictive of the DSGC’s PD (Figure 5 of Briggman et al., 2011). Spots represent synaptic inputs from SACs, which provide the direction-selective GABAergic inhibition to DSGCs. Each stick points in the direction of the SAC soma. This orientation also correlates with the stimulus direction in which minimal GABA will be released from that synapse. **D**, Magnified view of the highlighted area of the dendritic arbor in C (in blue) where SAC dendrites are less organized. If DSGCs integrated inputs globally (top) the average inhibition would be poorly tuned. This would result in all dendrites being similarly but weakly tuned for direction, as indicated by the length of the arrows at each site. However, if inputs were integrated locally (bottom), all dendritic branches in the region would inherit distinct tuning properties from SAC dendrites orientated in different directions.

It is now well-established that direction-selective GABAergic signals in DSGCs arise from the radiating dendrites of starburst amacrine cells (SACs; Taylor and Smith, 2012; Vaney et al., 2012; Wei, 2018). Since strong Ca^2+^ signals in distal SAC varicosities are only observed in response to centrifugal motion (from the soma to the dendritic tip) but not in response to motion in the opposite direction, the orientation of a given SAC dendrite provides an approximation of its directional preference (Euler et al., 2002; Morrie and Feller, 2018; Poleg-Polsky et al., 2018). In a seminal study using serial block-face electron microscopy, Briggman et al. (2011) demonstrated that SAC varicosities, where GABA release occurs, specifically contact DSGCs whose directional preference is opposite to their own (**Figure 1C**). This asymmetric wiring ensures that tuning properties of SACs are effectively translated into directionally tuned GABAergic inhibition, which plays a key role in shaping direction selectivity in DSGC dendrites. Thus, a close inspection of the anatomical connectivity between DSGC-SAC accurately predicts the DSGC’s directional preferences (Briggman et al., 2011).

Interestingly, the circuitry required for generating direction selectivity in ganglion cells appears to be distributed within small regions, ~50-100 µm diameter, over its receptive field (Barlow and Levick, 1965). Based on the natural branching patterns and the theoretical voltage length constants of ganglion cell dendrites, Koch et al. (1982) inferred that the DSGC dendritic arbor comprised of subunits similar in size to those suggested by Barlow and Levick. These subunits are electrically isolated from each other but relatively isopotential within and thus expected to compute direction independently. More recently, DSGC dendrites have been shown to support robust TTX-sensitive dendritic spiking, which enables information from distal parts of the dendritic tree to strongly influence somatic spike activity (Oesch et al., 2005; Sivyer and Williams, 2013; Trenholm et al., 2014). Together, these findings make a compelling case that DSGCs utilize a parallel processing scheme for computing direction.

However, how the excitatory and inhibitory inputs that shape direction selectivity are functionally organized at the level of individual dendrites has not been directly investigated, and is the main focus of this study. Cursory examination of the orientation of presynaptic SAC dendrites within sub-regions predicts that the directional tuning of GABAergic input would be heterogenous across the dendritic arbor (Briggman et al., 2011; **Figure 1C**). However, it must be noted that interpreting the wiring at a subcellular scale is difficult because there is significant variance in the directional tuning of individual varicosities on SAC dendrites with the same orientation, which is exacerbated by trial-to-trial variability (Ding et al., 2016; Morrie and Feller, 2018; Poleg-Polsky et al., 2018). Thus, while the macroscopic tuning properties of DSGCs are evident from the anatomical connectivity (Briggman et al., 2011), the nature of the directional information provided to individual dendrites is harder to predict. Determining the patterns of local inputs at the level of individual dendrites has significant implications for understanding how DSGCs accurately process motion information in parallel.

Previously, optical and electrophysiological methods have been employed to measure dendritic activity in DSGCs (Brombas et al., 2017; Sivyer and Williams, 2013; Yonehara et al., 2013), but the robust propagation of dendritic spikes obscures the contributions of local synaptic inputs to direction selectivity in dendrites. To circumvent this issue, in this study we used 2-photon imaging to measure dendritic Ca^2+^ signals from mouse ON-OFF DSGCs under conditions where voltage-gated sodium channels (NaV) were pharmacologically blocked. Under these conditions we characterize the local dendritic tuning properties, which allowed us for the first time to directly infer the degree to which the presynaptic excitatory and inhibitory inputs to DSGCs are functionally organized at a subcellular level.

## RESULTS

### Direction selectivity in dendrites in the absence of NaV channel activity

We reasoned that the degree to which individual dendrites are tuned for direction reflects the relative strength of the excitatory and inhibitory inputs over the region that they are spatially integrated. If E/I interact on broad spatial scales, then dendritic sections within relatively large regions should share their tuning and noise properties (**Figure 1D;** Koch et al., 1982; Schacter et al., 2010). However, if synaptic inputs interact more locally, directional responses should be independently formed within smaller dendritic segments. To distinguish these possibilities, we directly assessed the tuning properties of individual DSGC dendrites in a flat-mount retinal preparation, in which responses over large regions of their planar dendritic trees could be captured in single field of view using two-photon Ca^2+^ imaging. In these experiments, DSGCs were identified based on their spiking responses, and then filled with a Ca^2+^ indicator (200 µM OGB-1) through a patch electrode, which was also used to monitor the somatic voltage throughout the imaging session. Dendritic Ca^2+^ signals were recorded in response to spots moving in eight directions over the DSGC’s receptive field. To prevent dendritic spiking or somatic back-propagation from driving globally coordinated Ca^2+^ signals, the dendritic Ca^2+^ responses were measured in the presence of NaV channel blockers (2 mM QX-314 in the patch electrode, or 0.5 µM TTX in the bath solution; **Figure 1**).

Throughout the arbor, Ca^2+^ responses extracted from small regions of interest (3-4 µm^2^ ROIs) were well-tuned for the direction of motion. The preferred direction (PD; computed from the vector sum of responses evoked by spots moving in eight directions) of the dendritic Ca^2+^ signals closely matched that of the spiking response, which was recorded prior to the imaging session (**Figure 2B-E**). Although the average PDs of nearby dendritic sites were similar, they significantly varied over each stimulus set (8 directions per set), indicating that dendritic sites may be subject to different sources of noise. For example, comparing the directions encoded by four neighboring regions on the same dendritic branch (**Figure 2D**) it can be seen that on the first trial set, ROIs 3 and 4 deviate by ~37°, while ROIs 2 and 4 deviate by ~50°. On the next trial set, the deviations changed in sign and amplitude. Importantly, such deviations do not arise from technical noise sources (e.g. PMT shot noise), since the noise at each site before the stimulus onset was low compared to the peak amplitude of the evoked Ca^2^ transients (**Figure 2E**, grey shading). Note, for this and all subsequent analysis, only dendritic sites which responded to at least one direction of motion with a peak amplitude greater than three times the standard deviation of the baseline noise were considered. As we will discuss in a later section, larger variations in the light-evoked Ca^2+^ signals are likely attributable to the stochastic nature of local E/I interactions (**Figures 5–7**).

**Figure 2.**
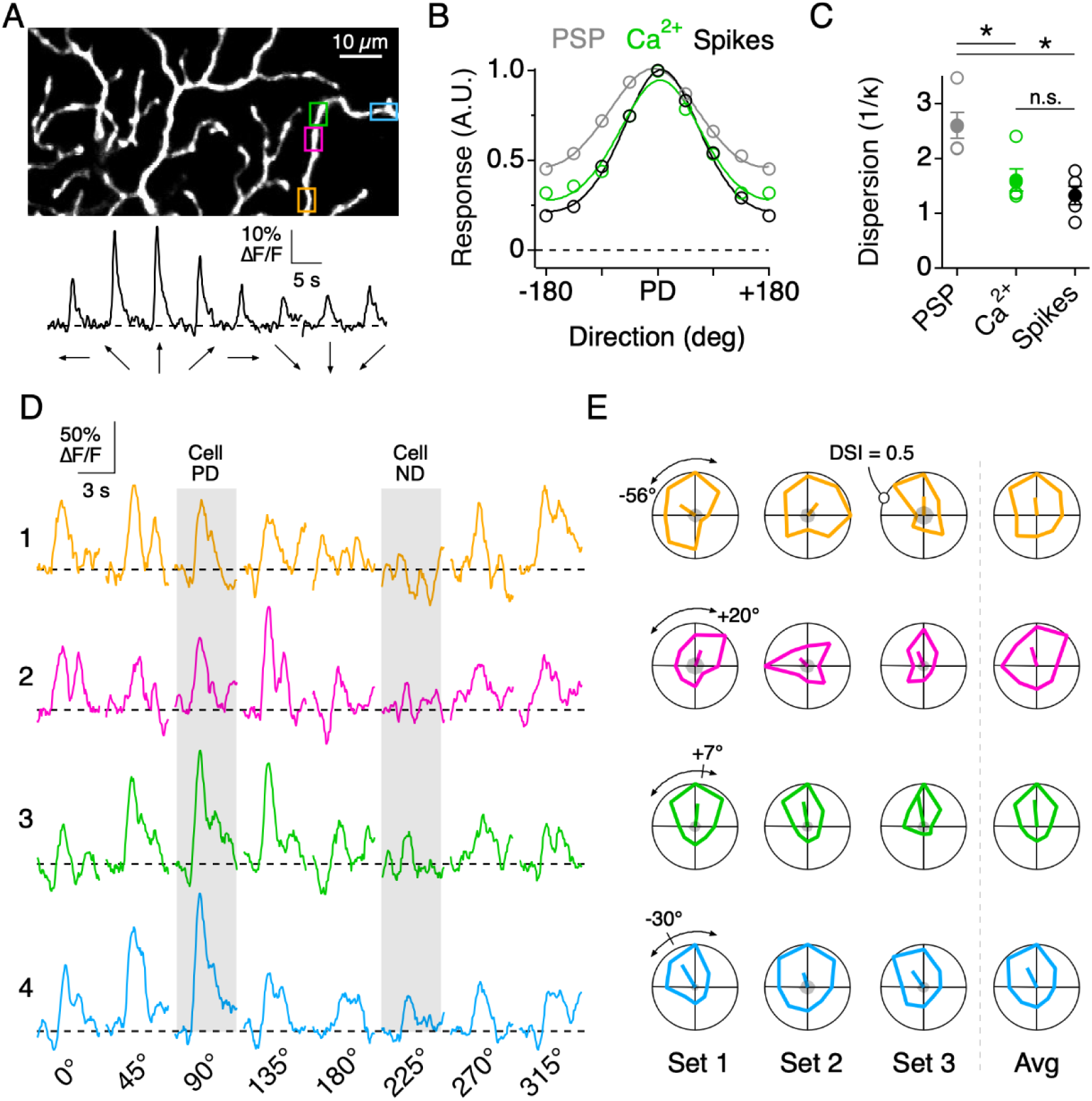
Directional tuning in DSGC dendrites in the absence of NaV-dependent spikes. **A**, Imaging area of the ON arbor of an ON-OFF DSGC. Bottom, Ca^2+^ signals over the entire imaging window in response to a positive contrast spot moving in 8 different directions. **B**, The average tuning curves of the spiking response, subthreshold somatic voltage (PSP), and dendritic Ca^2+^ signals (n = 7 cells). Solid lines indicate Von Mises fits to the data. The PD is normalized to that of the spiking response. **C**, The value 1/κ, extracted from the Von Mises fits, is a measure of tuning width, or angular dispersion, for circular data. The tuning curve from the somatic voltage was significantly broader than both the Ca^2+^ signals and the spiking responses. Asterisks indicate p < 0.05 (t test). **D**, Ca^2+^ signals over a single stimulus set (1 set = 8 directions) at four nearby ROIs on the same dendritic branch, as demarcated by the colored boxes in **A**. The PD and ND responses as measured by the spiking response are indicated by the shading. **E**, Polar plots of the peak Ca^2+^ responses over consecutive stimulus sets, with the PD of each dendritic ROI indicated by the line extending from the origin, the length of which denoting the DSI. The arcs above the polar plot indicate the range in the preferred angle across these four sites on a single stimulus set. Right, the average polar plot for each ROI over the displayed stimulus sets.

Similar to the single branch analyzed above, the direction encoded by dendritic sites distributed across larger fields of the arbor remained relatively constrained around the DSGC’s PD indicated by its spiking response. The direction encoded by a given site rarely fell in the wrong directional hemisphere, even when computed over a single set of trials (9/353 sites in 7 cells, 2.5%). The overall variation in the PD from single stimulus sets (σ_θ_ = 52.8°; 353 ROIs from 7 cells) was significantly reduced when three to five stimulus sets were averaged (σ_θ_ = 31.6°, **Figures 3A, C**). In addition, we found that the strength of the tuning, quantified as the direction selectivity index (DSI; see Methods), also varied among the population of dendritic sites (DSI = 0.19 ± 0.08; µ ± s.d.; 353 ROIs from 7 cells; **Figure 3B, D**). Interestingly, adjacent sites with similar PDs could have large differences in DSI, and vice versa, indicating that the directional tuning of each site arises relatively independently (**Figure 3E**).

**Figure 3.**
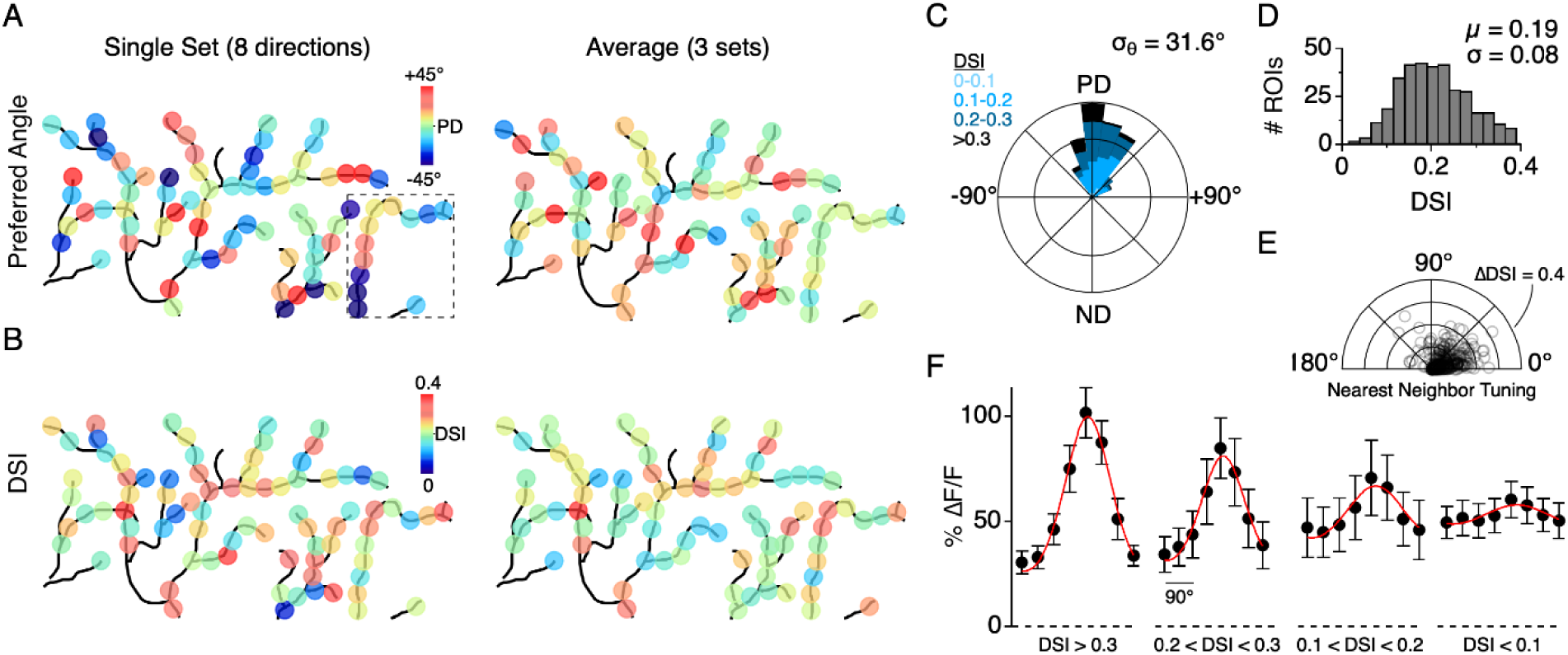
Distribution of directional tuning properties across the DSGC dendritic arbor. **A**, Maps of the preferred direction of each dendritic ROI, as measured over a single stimulus set (left) or averaged over multiple sets (right). Boxed area denotes the region used for analysis in **Figure 2. B**, Left, Same as in **A**, but for DSI. **C**, Rose plot showing the distribution of the average PD for ROIs across the dendritic arbors of 7 ON-OFF DSGCs, relative to the PD of each DSGC’s spiking response. Colors indicate the proportion of ROIs within different ranges of DSI. **D**, Top, Distribution of DSIs for all dendritic ROIs (DSI = 0.19 ± 0.08, n = 353 ROIs from 7 cells). **E**, Polar plot showing the difference in the DSI and PD between each ROI and its nearest neighboring ROI. **F**, Average tuning curve for dendritic ROIs within different ranges of DSI.

We also found that the peak magnitude of the Ca^2+^ signals varied as a function of the DSI (**Figure 3F;** 353 sites from 7 DSGCs). In response to preferred motion, sites with a low DSI (< 0.1) had significantly weaker Ca^2+^ responses than those with a high DSI (ΔF/F = 0.60 ± 0.09 versus 1.02 ± 0.12, p < 0.01, t-test, µ ± s.d.). However, the null responses at the low DSI sites were generally stronger than those observed at the high DSI sites (ΔF/F = 0.49 ± 0.08 versus 0.31 ± 0.05, p < 0.01, t-test, µ ± s.d.). Despite their weak tuning, sites with low DSI sites still computed the correct direction, deviating by 0.99 ± 52° from the PD computed from spiking (n = 37 sites in 7 cells, µ ± s.d.). Together these data suggest that presynaptic excitatory and inhibitory inputs are precisely organized, enabling a similar direction to be encoded in many points across the dendritic arbor.

It is important to note that nonlinearities inherent in the dendritic Ca^2+^-permeable channels are expected to produce signals that are more compartmentalized than the underlying local dendritic voltage (Wybo et al., 2019; Meier and Borst, 2019; Grimes et al., 2010; **Figure 7**). Interestingly, Ca^2+^ responses remained confined to small sections of the dendrite over several hundred milliseconds, indicative of the strong buffering mechanisms dendrites (Biess et al., 2011; Awatramani et al., 2007). Also, as these experiments were carried out in the presence of NaV blockers, it is possible that E/I interactions that take place over broad spatial scales are abolished, and the contribution of local synaptic inputs are artificially emphasized. However, the tuning of both the spiking and the dendritic Ca^2+^ responses (measured across the entire arbor) were similar to each other, but significantly narrower than that of the subthreshold somatic voltage (**Figure 2C**). This suggests that analogous or shared mechanisms for nonlinear amplification (e.g. voltage-dependent Ca^2+^ channels) may underlie the sharpening of the tuning curves of DSGCs, as previously described (Oesch et al., 2005). Thus, it is possible that the localized Ca^2+^ signals that we observed are computationally relevant (also see Discussion).

**Figure 7.**
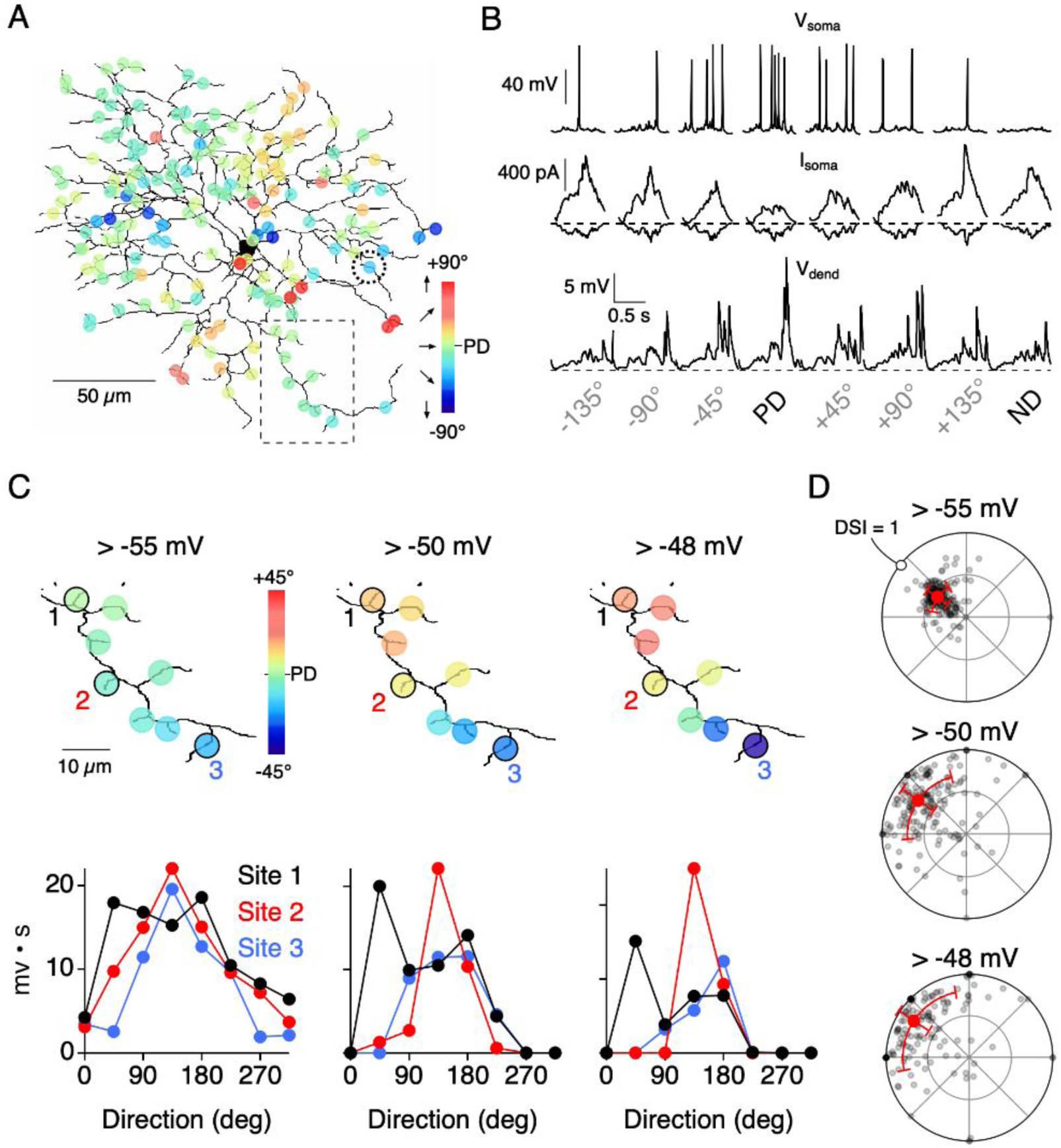
A computational model illustrating how threshold nonlinearities encourage dendritic response compartmentalization. **A**, Reconstruction of an ON-OFF DSGC used in a multicompartmental NEURON (Poleg-Polsky and Diamond, 2016; see text for details), illustrating how stochastic transmission can give rise to different responses in nearby dendritic sites in the absence of NaV. **B**, Spiking (V_soma_), excitatory and inhibitory currents (I_soma_) measured from the soma; and voltage measured from a single dendritic site (V_dendrite_; from site circled in A) in response to a simulated edge moving across the receptive field in 8 different directions. The angle of the vector sum of the integrated voltage responses (above a threshold of −55 mV) was used to compute the PDs. **C**, Expanded view of the dendritic branch indicated by the boxed area in **A**, showing how the stringency of the voltage threshold (indicated above images) can affect the directional tuning in nearby dendritic sites. The integrated voltage at three sites (Site 1-3) is shown below. **D**, Polar plots showing the distribution of PDs and DSIs measured over the dendritic tree, measured using different threshold values. Note, in the model all sites have the same average tuning, and the illustrated differences in tuning between sites only occur on single trial sets (8 directions) due to the stochastic nature of transmitter release.

### The role of NMDA and GABA receptors in shaping directional tuning of dendrites

In DSGCs, glutamatergic inputs are detected in part by NMDA receptors (Poleg-Polsky and Diamond, 2016; Sethuramanujam et al., 2017), which also serve as a potential source of Ca^2+^ influx. Thus, the observed variance in the dendritic Ca^2+^ responses could in theory reflect variability in presynaptic glutamate release. However, previous work suggests that the voltage-dependent properties of NMDA receptors allow them to act ‘multiplicatively’ on the DSGC membrane potential. That is to say, NMDA receptors amplify responses without changing the directional tuning properties (Poleg-Polsky and Diamond, 2016; Sethuramanujam et al., 2017). Thus, the activation of NMDA receptors does not appear to change the E/I balance that underlies direction selectivity in DSGCs. Consistent with this notion, we found that Ca^2+^ responses measured in the presence of the NMDA receptor antagonist D-AP5 (50 µM) were weaker but still well-tuned for direction (**Figure 4A**). Under these conditions, in which Ca^2+^ responses are expected to be reliant mainly on voltage-gated Ca^2+^ channels, the distribution of tuning angles (p = 0.49, Angular Distance test; T [0.55] < TC [1.96], Watson-Williams test; **Figure 4C**) and DSIs (p = 0.15, Kolmogorov-Smirnov test, **Figure 4D**) throughout the dendritic arbor were indistinguishable from that measured under control conditions. Thus, NMDA receptors appear to multiplicatively scale dendritic Ca^2+^ signals as they do the somatic voltage signals (Poleg-Polsky and Diamond, 2016; Sethuramanujam et al., 2017). This unique property makes NMDA receptor activity particularly useful for visualizing E/I balance in the dendritic tree without disrupting it.

**Figure 4.**
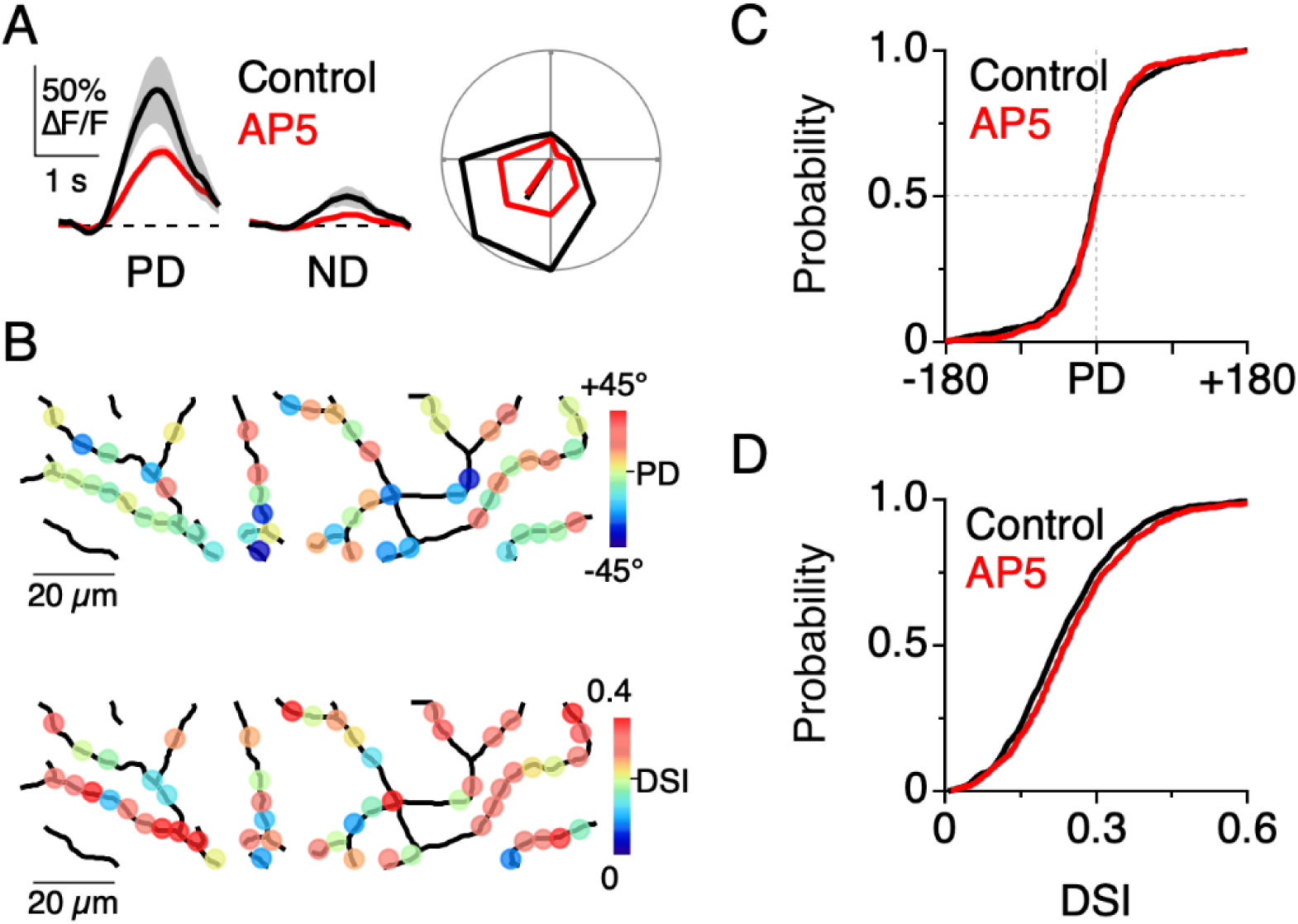
NMDA receptors amplify dendritic responses without altering direction selectivity. **A**, Example Ca^2+^ signals in response to preferred and null motion in control conditions (black) and in the presence of the NMDA receptor antagonist, D-AP5 (50 µM; red). Shading is ± standard error. Right, Example polar plot showing the directional tuning of dendritic Ca^2+^ signals before and after NMDA receptor blockade. **B**, Map of the PD (top) and DSI (bottom) for dendritic ROIs during NMDA receptor blockade. **C**, Cumulative distributions of the PD for dendritic ROIs. **D**, Cumulative distributions of the DSI for dendritic ROIs.

In contrast to blocking NMDA receptors, reducing SAC-mediated GABAergic inhibition had a profound effect on the dendritic tuning properties. Selective disruption of GABA release from SACs was achieved by using the conditional deletion of the vesicular GABA transporter, which strongly reduces GABA release from SACs (vGAT KO^fl/fl^∷ChAT^Cre^; Pei et al., 2015; Blenkishop et al., 2019). In these transgenic lines directional tuning is heterogeneously compromised, but we selected DSGCs with poor tuning properties (DSI < 0.10), which signified that inhibition was reduced. Not surprisingly, the directional tuning of dendritic Ca^2+^ signals were significantly compromised in these knock out mice (DSI = 0.06 ± 0.03, µ ± s.d., n = 4; **Figure 5A-C**), similar to the spiking responses measured prior to the imaging session. These results are consistent with the idea that the combined excitatory signals mediated by glutamate and acetylcholine receptors are not strongly tuned for direction (Park et al., 2014; Yonehara et al., 2013; but see Pei et al., 2015). Thus, the direction selectivity observed in wild type DSGC dendrites must be shaped by directionally tuned inhibition, as originally envisioned (Barlow and Levick, 1965).

**Figure 5.**
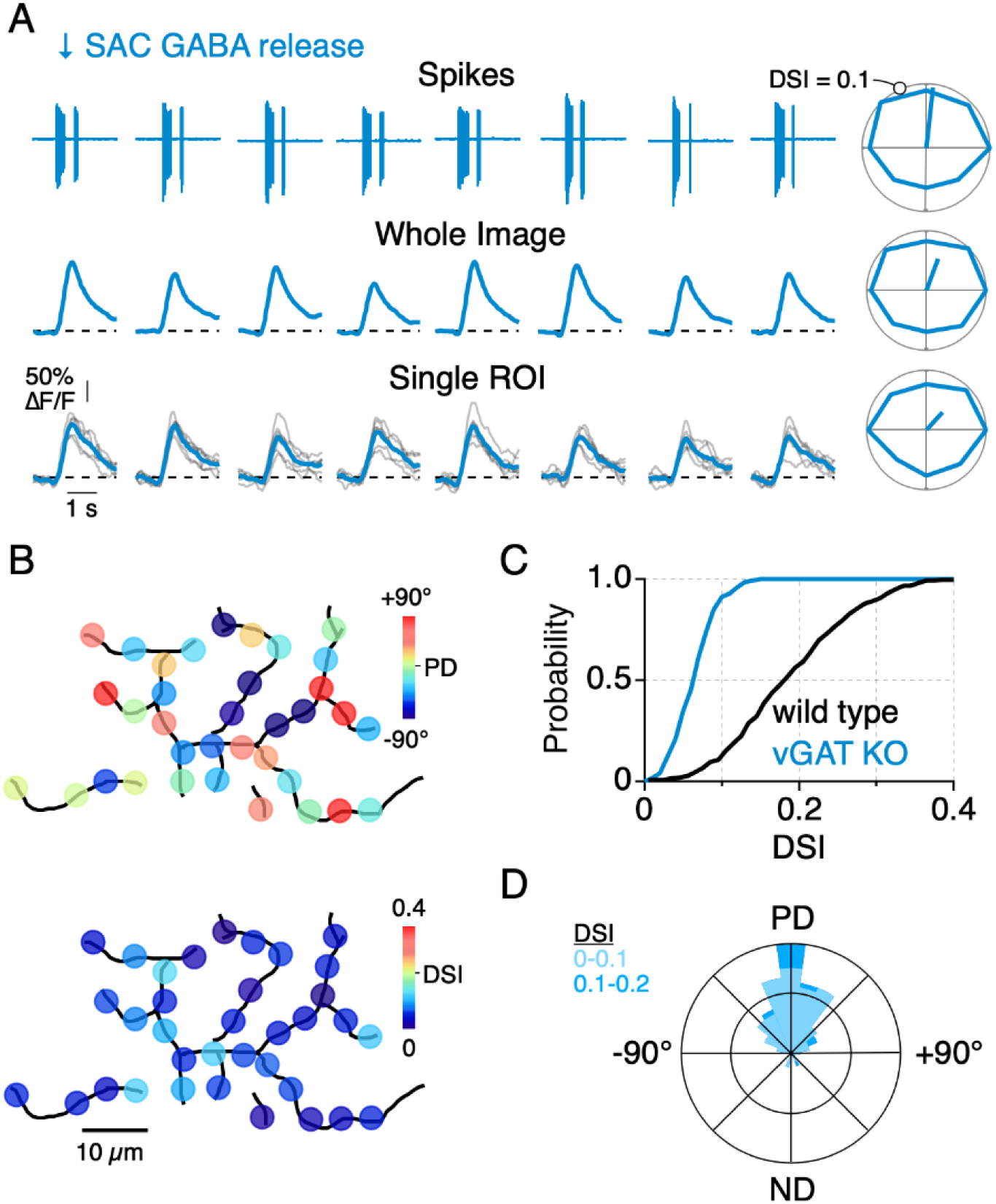
Dendritic Ca^2+^ signals are tuned by SAC inhibition. **A**, Spiking response and Ca^2+^ signals from a genetically labelled ON-OFF DSGC. Right, Polar plots of the spike rate and peak Ca^2+^ responses. Lines extending from the origin indicate the preferred direction and the DSI (note, DSI scale is from 0–0.1). **B**, Maps of the PD (top) and DSI (bottom) at dendritic ROIs in vGAT-KO mice. **C**, Cumulative distribution of the DSI from wild type and vGAT KO mice. **D**, Rose plot of the preferred direction of dendritic ROIs in vGAT-KO mice. Colors indicate the proportion of ROIs within different ranges of DSI.

Measurements in the vGAT KO mice also highlight the sensitivity of the Ca^2+^ imaging method for detecting changes in E/I. Specifically, subtle asymmetries present in the spiking responses, which would usually be considered ‘non-directional’ (Pei et al., 2015), were also present in the dendritic Ca^2+^ responses in these mice (**Figure 5B-D**). Although the dendritic Ca^2+^ signals extracted from single ROIs in the dendritic tree were weakly tuned for direction, they encoded a similar direction to the spiking response, indicating that the weak tuning did not arise from technical noise sources (**Figure 5B, D**). These data also support our initial claim that tuning of the dendritic Ca^2+^ responses closely reflect that of the somatic spiking (**Figure 2B-C**). The close correspondence in the tuning of the spiking and the dendritic Ca^2+^ responses (measured in the absence of NaV-dependent spikes) suggest that they are intimately linked.

### Independent synaptic processing within small dendritic segments

Next, to quantitatively assess the independence of nearby dendritic sites, we examined how Ca^2+^ signals at pairs of ROIs co-varied as a function of the cable distance between them. For this analysis, Ca^2+^ signals were measured over at least 30 consecutive presentations of orthogonal motion, where trial-to-trial variance is high (**Figure 6G**). The raw Ca^2+^ signals from each ROI were cross-correlated with signals from every other ROI in the imaging window. ROIs were found to be strongly correlated regardless of their distance of separation, since Ca^2+^ signals arise at all sites during the period of stimulus presentation (**Figure 6,Supplemental Figure 1**). However, when the mean-subtracted ‘noise’ residuals measured from each ROI were cross-correlated, only nearby sites were significantly correlated with each other (**Figure 6C-E**). Noise correlations measured in neighboring ROIs were not present when trials were shuffled (**Figure 6C**). The correlation strength rapidly decayed over distance with a space constant of λ = 5.3 µm (**Figure 6D-E**; n = 6 cells), suggesting that Ca^2+^ signals beyond this distance arise independently. Given the dense spacing of excitatory and inhibitory synapses in DSGC dendrites (Bleckert et al., 2013; Briggman et al., 2011; Jeon et al., 2002; Sigal et al., 2015), the strong compartmentalization over 5-10 µm dendritic segments suggests that Ca^2+^ signals are most strongly influenced by only a few synapses.

**Figure 6.**
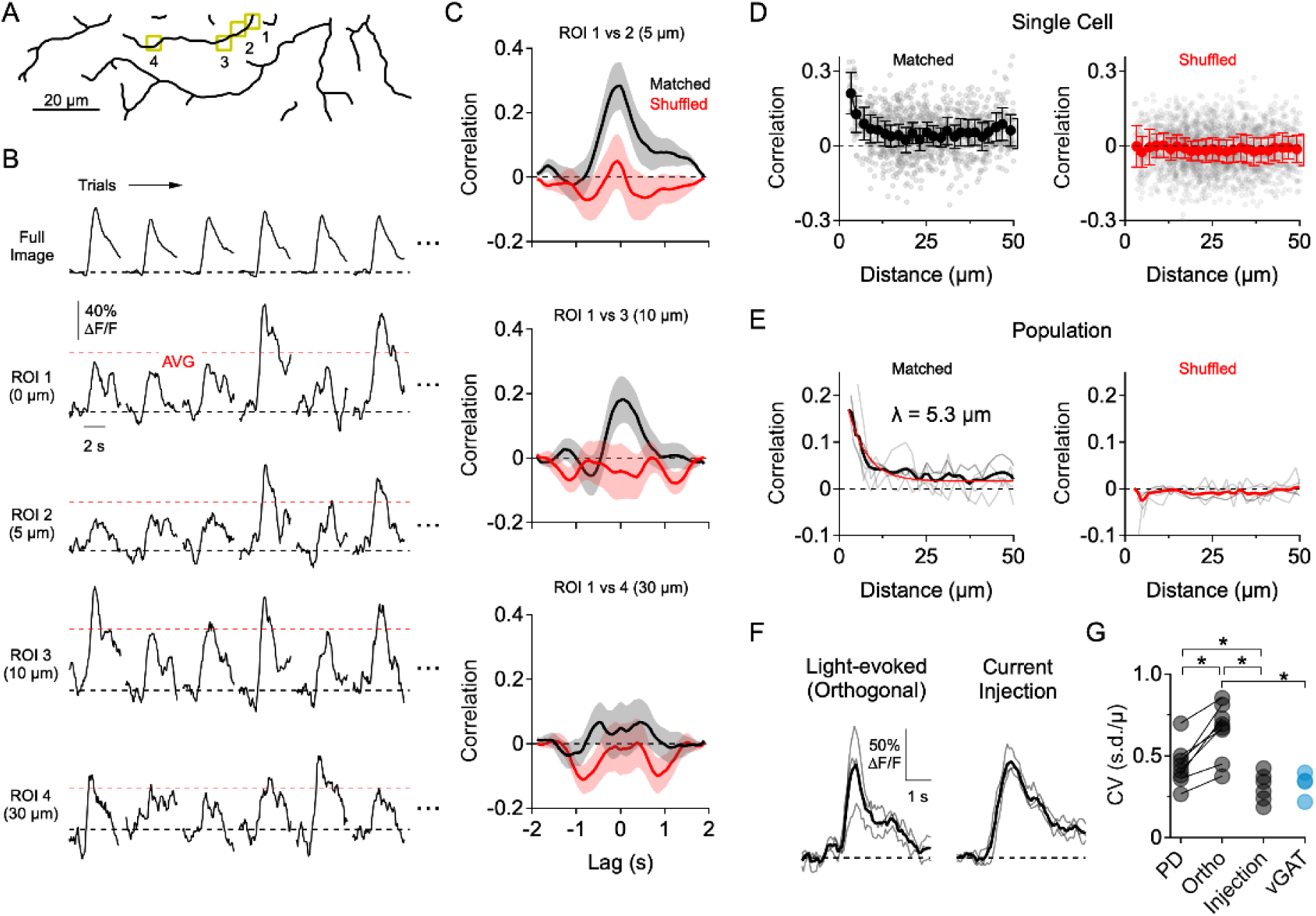
Optical fluctuation analysis reveals the spatial extent to which dendritic Ca^2+^ signals form independently. **A**, Reconstruction of the imaging area in the ON arbor of an ON-OFF DSGC. **B**, Ca^2+^ signals over 30 consecutive trials in response to motion in a direction orthogonal to the preferred-null axis (only 6 trials are shown). Ca^2+^ signals are from the ROIs highlighted in **A**, or from the entire arbor within the imaging window (Full Image). Red dotted lines indicate the average peak Ca^2+^ signal over all trials. **C**, Cross-correlation of the mean-subtracted residual Ca^2+^ signals between ROI 1 and the other highlighted ROIs, each of which is at an increased distance. Correlations were performed using matched trials (black) or shuffled trials (red). **D**, Peak correlation versus distance between the ROIs for matched (black) and shuffled trials (red) in a single DSGC. Faded data points are individual correlation values, solid points are the average (±s.e.m.) over 2 µm bins. **E**, Peak correlation versus distance for a population of 7 ON-OFF DSGCs for matched and shuffled trials. Faded lines are the average correlation versus distance data for single cells, bold lines are the population average. For matched trials, the thin red line is an exponential fit, with a space constant of λ = 5.3 µm. **F**, Light-evoked Ca^2+^ signals from an example ROI in response to orthogonal motion compared to Ca^2+^ signals evoked by current injection at the soma. Thin lines are individual trials; thick lines are the average. **G**, Coefficient of variance (standard deviation divided by the mean) for the peak Ca^2+^ signals evoked by preferred motion, orthogonal motion, somatic current injection, and in response to motion in a mouse that has the vesicular GABA transporter knocked out of SACs (vGAT^fl/fl^∷ChAT^Cre^). Asterisks indicate p < 0.05, Student’s t-test.

Several pieces of evidence suggest that the trial-to-trial variability in the light-evoked Ca^2+^ signals arise primarily from variations in inhibitory synaptic activity. First, trial-to-trial variability was significantly lower when Ca^2+^ signals were evoked by directly depolarizing the DSGC through the patch electrode in the presence of synaptic blockers (50 µM DL-AP4, 20 µM CNQX, 10 µM UBP-310; **Figure 6F-G**). These data also suggest that voltage-dependent Ca^2+^ channels are a major source of Ca^2+^ influx in the presence of 2 mM QX-314 (note, low concentrations were used because at 10 mM QX-314 reduces Ca^2+^ currents; Talbot and Sayer, 1996). Second, the response variability during orthogonal motion was significantly reduced in the vGAT KO compared to wild type mice, in which SAC inhibition is intact (**Figure 6G**). Third, in wild type mice, the response variability to motion in the PD—in which inhibition is quite low—was significantly less than that of orthogonal motion. Overall, the results support the notion that the fidelity of excitatory transmission is high, and that the increased variability of orthogonal responses is due to postsynaptic E/I interactions. Since inhibition helps compartmentalize dendritic responses (Lovett-Barron et al., 2012; Poleg-Polsky et al., 2018), it is possible that part of the decrease in variability under conditions with lower GABAergic input results from integrating excitatory inputs over a larger area. However, the low trial-to-trial variance of the Ca^2+^ signals in the PD/vGAT KO made it difficult to perform the same fluctuation analysis as was done for orthogonal motion. Instead, examining the latencies of responses at higher temporal resolution revealed a sequential activation of individual dendrites as a preferred moving stimulus traversed the DSGC’s receptive field (**Figure 6, Supplementary Figure 2**). Although the precise spatial dimensions of E/I interactions could not be established for these conditions, this result indicates that individual dendritic branches function relatively independently, even under conditions where there is low inhibition.

Similarly, in response to motion in the null direction, in which GABAergic inhibition is strong, we could measure dendritic activity in isolated dendritic ‘hot spots’ (FWHM 3.0 ± 1.2 µm; n = 61 sites in 7 cells; **Figure 6, Supplemental Figure 3**). Null signals at a given site were not consistently observed over multiple trials, suggesting that they arise from probabilistic transmission (failure of inhibition/excess excitation) rather than from the absence of inhibitory synapses in that dendritic region. Since inhibitory conductances are largest during null direction motion, the hot spots measured under this condition likely represent the highest degree of response compartmentalization in DSGC dendrites.

### Dendritic nonlinearities promote dendritic independence

Synaptically-evoked Ca^2+^ signals are expected to be more compartmentalized than dendritic voltage (Wybo et al., 2019; Meier and Borst, 2019; Grimes et al., 2010), since they arise from inherently nonlinear sources (voltage-gated channels and NMDA receptors). Such nonlinearities are also expected to promote the apparent independence of dendritic activity (Grimes et al., 2010), and could explain how nearby dendritic regions could express different directional tuning properties. To further test this idea, we constructed a multi-compartmental computational model of a reconstructed ON-OFF DSGC (**Figure 7A**; Poleg-Polsky and Diamond, 2016) driven by 177 inhibitory and excitatory inputs, which summed to produce currents similar to those measured experimentally at the soma (in ‘voltage-clamp’ mode), and evoked robust direction selective spiking output (**Figure 7B)**. The inhibitory input at each site was directionally tuned in an identical manner, while the excitatory inputs were untuned. The changes in membrane voltage in response to a simulated edge moving across the dendritic arbor in different directions was recorded at each site.

Since neurotransmitter release is stochastic, the precise E/I balance for a given direction varied between sites and from trial-to-trial. Thus, the resulting directional tuning of local dendritic voltage signals (measured in the absence of NaV) differed between neighboring sites, even though all sites had similar average tuning properties. Similar to the Ca^2+^ imaging data, the angle encoded at most sites varied about the DSGC’s PD (**Figure 7B, C**). Since Ca^2+^ signals rely on the activation of CaV channels, which become steeply dependent on voltage above −55 mV (Randall and Tsien, 1997), we examined how dendritic tuning varies when responses are thresholded in this range. We found that systematically thresholding the voltage data from −55 mV to −48 mV led to an increase in both the angular variance (σ_θ_ = 25.4°, 40.4°, and 45.0° for −55 mV, −50 mV, and −48 mV thresholds, respectively) and the DSI (DSI = 0.42, 0.70, and 0.80 for −55 mV, −50 mV, and −48 mV thresholds, respectively; **Figure 7D**). When responses were strongly thresholded, even sites located on neighboring branches encoded different directions, indicating their functional independence (**Figure 7C**). Together, the model data support the idea that dendritic nonlinearities could help promote compartmentalized direction tuning in dendrites.

### SAC input exerts local control of dendritic direction selectivity

So far, the major inference that we can draw from our results is that the directional tuning of small dendritic sections is shaped primarily by local inhibitory synaptic inputs. As a final proof of principle, we examined dendritic tuning after locally disrupting inhibitory inputs arising from a limited set of SACs. To do so we initially tried local puffs of GABA receptor antagonists to block inhibition onto specific dendrites but found it difficult to keep the drug sufficiently localized. We also tried controlling GABA release from distal SAC varicosities through a somatic patch electrode but found that this was not possible in the presence of light-evoked synaptic conductances. Finally, we resorted to single cell ablation methods using sharp electrodes to disrupt a some of the inhibitory input mediated by SACs (Jacoby et al., 2015). In a mouse line in which SACs were genetically labelled (ChAT-cre∷ nGFP mouse line; Vlasits et al., 2014), 3-7 SACs with somas located on the null side of the DSGC’s receptive field—which likely provide multiple strong GABAergic contacts to the DSGC—were targeted for ablation (**Figure 8A**). SAC ablation increased the spiking activity in the null direction and thus diminished direction selectivity, confirming that ablation of a few SACs results in a decrease in null inhibition (**Figure 8B**).

**Figure 8.**
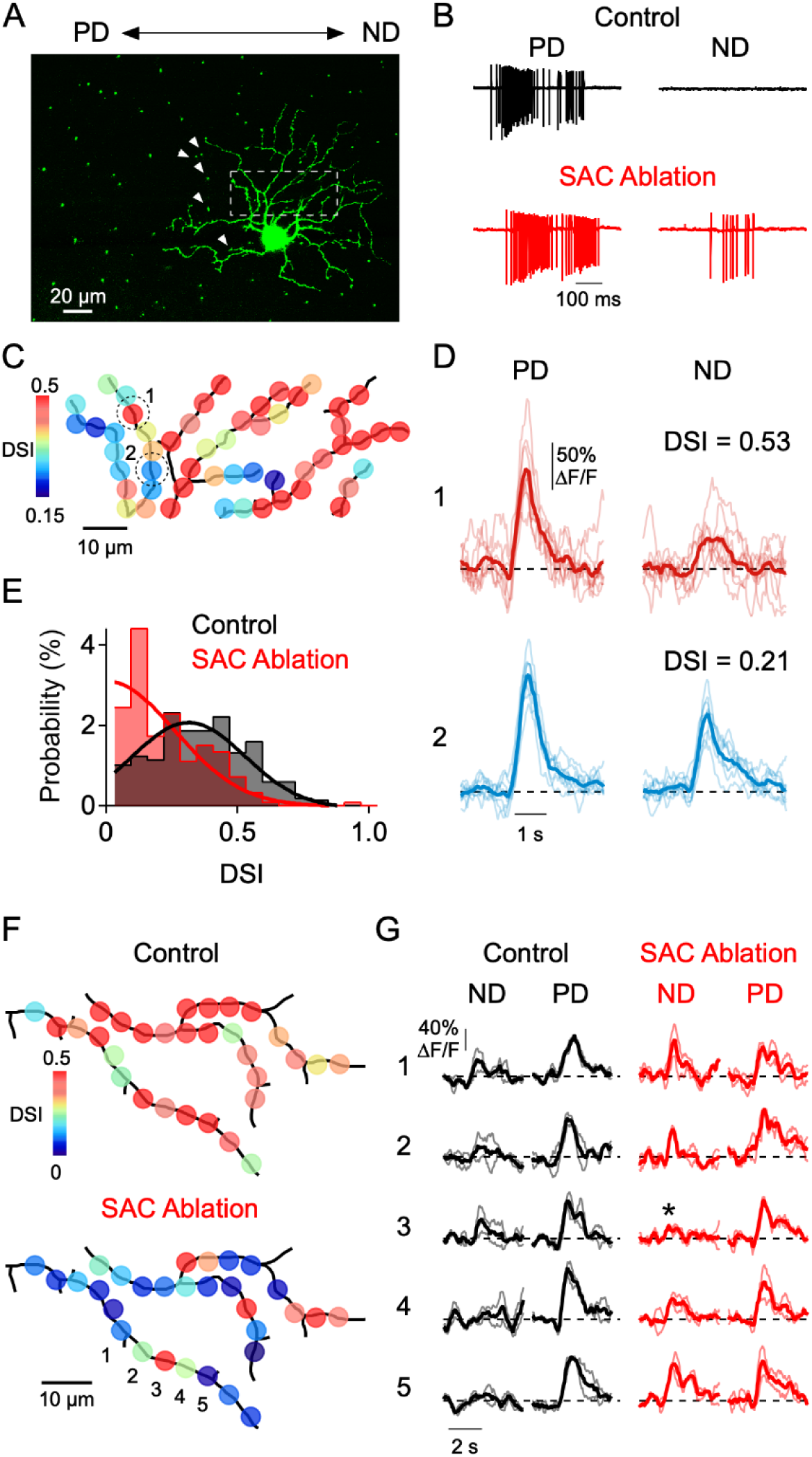
Spatially restricted disruption dendritic direction selectivity by restricted null-SAC ablations. **A**, Image showing the mosaic of SAC somas labelled with a nuclear GFP marker surrounding a DSGC filled with OGB-1. Arrowheads indicate null-side SACs that were targeted for electrical ablation with a micro electrode. **B**, Spiking responses of a DSGC before (black) and after SAC ablation (red). **C**, DSI map of the imaged area (dotted square in A), after 5 SACs were ablated. **D**, Average Ca^2+^ signals (10 trials) in 2 ROIs (indicated in **C)** with strong and weak DSI. **E**, Distribution of DSIs for a population of sites in 4 DSGCs after SAC ablation (red), compared with the distribution from DSGCs imaged under control conditions (black). Note, DSI was calculated from only preferred and null stimuli, and are generally of higher values than DSIs reported in Figures 3–4 computed from 8 directions. Solid lines are Gaussian fits to the data. **F**, Maps of the DSI in a DSGC before and after SAC ablation. **G**, Average Ca^2+^ signals from five ROIs (1-5) in F, before (black) and after (red) SAC ablation. Asterisk denotes a strongly tuned dendritic site flanked by poorly tuned regions.

Notably, after SAC ablation we found that the directional tuning was highly heterogeneous within imaged regions of the DSGC dendritic arbor. Many dendritic sites were well-tuned for direction, similar to control conditions, while others were poorly tuned (**Figure 8C, E**). To abbreviate these otherwise lengthy experiments, stimuli were only moved along the preferred-null axis. The overall DSI distribution was significantly different than that measured without disrupting SAC release (p = 10^−16^, Kolmogorov-Smirnov test; **Figure 8F**). Consistent with the idea that SAC ablation results in a reduction in inhibition, the null direction Ca^2+^ signals at poorly tuned sites were relatively large (**Figure 8D**). This contrasts with control conditions, in which poorly tuned sites had weak null and preferred responses (**Figure 3F**). Importantly, neighboring dendritic sites could have dramatically different range of DSI (**Figure 8D**). In some cases, we were able to measure the DSI before and after SAC ablation. Once again, clear changes in tuning strength occurred at many, but not all, sites throughout the arbor (**Figure 8G**). Sites that were unaffected by SAC ablation were closely interspersed among strongly affected sites, suggesting that the inhibitory inputs to a given site have little influence on the tuning of its neighbors (**Figure 8H**). These data convincingly demonstrate that local excitatory and inhibitory input shape direction selective Ca^2+^ responses in small dendritic sections.

## DISCUSSION

Since the theoretical studies of Rall (1962), there has been broad interest in understanding the role of dendrites in neural processing (Stuart and Spruston, 2015). Being the receiving units of neural information, dendrites are well positioned to perform complex computations on their inputs that are critical for behavior (London and Häusser, 2005). Several recent in vitro studies have applied innovative, but artificial, stimulation methodologies to show that E/I interactions occur over a fine spatiotemporal scale (within tens of microns/milliseconds; Hao et al., 2009; Liu, 2004; Lowe, 2002; Marlin and Carter, 2014; Müller et al., 2012; Müllner et al., 2015; Takahashi et al., 2016), a notion that has also been reinforced by modeling studies (Doron et al., 2017; Jadi et al., 2012). Here, we show that in the DSGC circuit, the natural patterns of presynaptic activity are coordinated with enough precision to support computations based on fine scale E/I integration.

### Presynaptic coordination of E/I

In the DSGC circuit, the degree of direction selectivity in dendrites is directly related to the relative strength of local E/I, since selectively blocking SAC inhibition renders dendritic Ca^2+^ responses non-directional (**Figure 5**). It must be stressed, however, that feature selectivity alone does not inform us about the local organization of excitatory and inhibitory inputs. For example, functional *in vivo* imaging studies have observed orientation-selective responses within small dendritic compartments (Jia et al., 2010; Wilson et al., 2016). In these cases the dendritic feature selectivity appears to be inherited from already-tuned excitatory inputs (Wertz et al., 2015), and it’s unclear whether postsynaptic interactions between E/I are involved.

For DSGCs, the notion that direction-selective responses arise from the integration of distinct patterns of E/I was expected based on many previous studies (Hanson et al., 2019; Vaney et al., 2012; Wei, 2018). However, the novelty of the current work lies in demonstrating that E/I drive accurate direction-selective information independently within single dendritic branches throughout the arbor. This conclusion is drawn from two major findings. First, manipulating sources of excitation (NMDA receptors; **Figure 4**) and inhibition (GABA receptors; **Figure 5**) affected the directional tuning of dendritic Ca^2+^ signals in predictable ways, suggesting that directional tuning relies on E/I interactions. Second, covariation analysis showed that dendritic Ca^2+^ signals arise independently at dendritic sites > 5-10 µm apart, suggesting that the underlying excitatory and inhibitory inputs interact independently over short dendritic lengths (**Figure 6**). Consistent with this notion, differences in the tuning properties (observed on single sets) were measured at nearby dendritic sites on the same dendritic branch (**Figures 2–3**). Finally, ablating a few SACs profoundly disrupts directional tuning at some DSGC dendritic sites, but not at other neighboring sites on the same dendritic branch (**Figure 8**), directly demonstrating that local inhibitory inputs shape directional tuning under the current experimental conditions. The impressive accuracy and homogeneity of directional tuning across the dendritic arbor suggests that the underlying excitatory and inhibitory inputs are highly coordinated throughout the DSGC’s dendritic tree.

Anatomical studies demonstrate that excitatory and inhibitory synapses are spaced at ~1-2 µm intervals along the dendrites of DSGCs (Bleckert et al., 2013; Briggman et al., 2011; Jeon et al., 2002; Sigal et al., 2015), indicating that the appropriate levels of E/I shaping direction selectivity are provided by only a few, high fidelity synapses. Indeed, SAC varicosities make large ‘wrap-around’ synapses with large pools of vesicles (Briggman et al., 2011), which could provide reliable GABAergic inhibition as well as cholinergic excitation. In addition, glutamatergic inputs are mediated by bipolar cell ribbon synapses, which can release multiple vesicles in rapid succession (James et al., 2019). These unique specializations may underlie the high fidelity of excitation and inhibition that endow DSGC dendrites with robust direction selectivity over such restricted spatial domains.

### Functional versus anatomical characterization of the circuit

The distribution of directional tuning that we observed is more homogeneous than expected given the anatomical and functional variance of presynaptic SACs (Briggman et al., 2011; Poleg-Polsky et al., 2018). We did not observe systematic shifts in the tuning across different dendritic sectors, as suggested by the anatomy (**Figure 1**), and rarely observed sites that were tuned to the wrong directional hemisphere (**Figure 2–3**). The directional tuning of dendritic sites was relatively homogeneous; 95% of the dendritic sites fell within 63° of the DSGC’s true PD (two standard deviations). This is a much tighter distribution than can be estimated from the anatomical connectivity, which shows SAC dendrite orientations roughly equally represented across a 90° spread (Briggman et al., 2011). However, predicting the precise directional preference of SAC dendrites from their anatomical orientation is a coarse and often inaccurate approximation, as demonstrated by recent studies (Ding et al., 2016; Morrie and Feller, 2018; Poleg-Polsky et al., 2018). It is also possible that in our study, sites with weak Ca^2+^ responses reflect some of the miswiring, which could lead to strong inhibition in both the null and preferred directions. Regardless of the detailed cellular mechanisms, the DSGC circuit appears to be better functionally organized at the subcellular level than previously envisioned.

### Implications for local dendritic integration

The fine-scale functional organization of the presynaptic circuitry implies that DSGCs may be capable of highly parallel processing of motion information. Modeling studies suggest that nonlinear integration can occur independently in highly branched dendritic structures (Koch et al., 1982; Rall, 1964; Wybo et al., 2019). Moreover, local inhibitory input dramatically increase the ability of neighboring sites to function as independent units (Wybo et al., 2019), as we have observed for DSGCs. Although our data cannot speak to whether each site is independently ‘integrating’ synaptic input in the sense of triggering a Na^+^ spike, we can say that direction-selective information is present at each site, which could certainly support local spike generation with high directional accuracy. If this is the case, then a compelling prediction is that small integrative subunits would allow DSGCs to constantly resample motion information over their receptive fields, permitting fast adjustments in spiking to new or changing directional information.

In our study, indirect evidence for local integration comes from examining null responses, where activity is highly localized to a few distributed hot spots. Since low levels of somatic spiking often occur in response to null motion, the dendritic spikes from which they originate are most likely generated over a small region of the dendrite. Further supporting this idea, paired recording studies in rabbit retina show that dendritic spiking in DSGCs can be triggered by the cholinergic synapses arising from a single SAC, which may only contact a few DSGC dendrites (Brombas et al., 2017). Similarly, during strong visual stimulation, dendritic action potentials initiated from multiple independent points in the dendritic tree may act cooperatively to increase the probability of spike initiation in a common parent dendrite (Sivyer and Williams, 2013). This scheme requires E/I to be appropriately coordinated at every point of the dendritic tree, as we have documented here. Otherwise, misplaced or mistimed conductances are unlikely to interact, and small imbalances in E/I would tend to be amplified by active dendrites, which could be detrimental to the neuron’s output (Larkum and Nevian, 2008; Liu, 2004).

The local dendritic integration that our data supports contrasts with global integration schemes that have also been implicated in direction encoding. Theoretical work suggests larger integrative subunits, on the order of 50-100 µm of dendrite (Koch et al., 1982; Schacter et al., 2010). Furthermore, a recent analysis of subcellular glutamate transients in ON DSGC dendrites suggests that the glutamatergic excitatory input comes from kinetically distinct bipolar cell types (Matsumoto et al., 2019). Here, optimal temporal summation during preferred motion occurs over the course of the DSGC’s entire dendritic arbor. Further experimentation is required to determine whether global integrative mechanisms operate under different visual scenarios, or whether they work together with local mechanisms to reinforce decisions being made by individual dendrites.

## Conclusions

Overall, the present study gains a deeper understanding of the local interactions of excitatory and inhibitory inputs in dendrites, and how they shape neural computations. Previous work has discussed single dendrites as being the basic computational units of the brain (Branco and Häusser, 2011, 2010). Our data suggest that during natural patterns of activity, accurate direction-selective information is present within small sections of the dendrites, raising the intriguing possibility that single dendrites may process motion information in parallel. Interestingly, recent evidence suggests that direction-selective neurons in the visual cortex also rely on E/I interactions (Wilson et al., 2018), and their circuits may be organized with more local specificity than previously envisioned (Scholl et al., 2017). Whether single neurons in cortical circuits have similarly tight spatial and temporal E/I coordination in order to prevent errant dendritic spiking remains to be determined. The recent advent of tools to optically monitor dendritic excitation and inhibition (iGluSnFR (Marvin et al., 2013) and iGABASnFR (Marvin et al., 2019) paves the way for exciting future investigations of how inhibition and excitation shape direction selectivity and other neural computations carried out by diverse circuits in the brain.

## METHODS

### Animals

Experiments were performed using adult (P30 or older) mice of either sex: C57BL/6J (JAX:000664). SACs were genetically accessed using the choline acetyltransferase (ChaT) Cre-mouse line (B6.129S6-ChATtm1(cre)lowl/J: JAX 006410). Cre-dependent expression of nuclear-localized GFP in SACs was achieved by crossing the ChAT-Cre with B6.129-Gt (ROSA)26Sortm1Joe/J (JAX: 008516). To reduce SAC GABA release the ChaT-Cre line was crossed with floxed Slc32a1tm1Lowl (also commonly referred to as Vgatflox/flox; JAX 012897). To target DSGCs the Trhr-EGFP (kindly provided by Dr. Marla Feller, UC Berkeley), Hb9-EGFP (JAX) lines were used in combination. Animals were housed in 12-hour light/dark cycles, in up to 5 animals per cage. All procedures were performed in accordance with the Canadian Council on Animal Care and approved by the University of Victoria’s Animal Care Committee.

### Tissue preparation

Mice were dark-adapted for at least 45 minutes before being anesthetized with isoflurane and decapitated. Retinas of both eyes were extracted in standard Ringer’s solution under a dissecting microscope equipped with infrared optics. Isolated retinas were laid flat, ganglion cell side up, over a pre-cut window in 0.22 mm membrane filter paper (Millipore). Mounted retinas were placed into a recording chamber and perfused with Ringer’s solution (110 mM NaCl, 2.5 mM KCl, 1mM CaCl_2_, 1.6 mM MgCl_2_, 10 mM glucose, 22 mM NaHCO_3_) heated to 35°C and bubbled with 95% CO_2_/5% O_2_. Retinas were illuminated with infrared light and visualized with a Spot RT3 CCD camera (Diagnostic Instruments) through a 40x water-immersion objective on a BX-51 WI microscope (Olympus Canada).

### Visual Stimulation

Visual stimuli were produced using a digital light projector and were focused onto the photoreceptor layer of the retina through the sub-stage condenser. The background luminance was measured to be 10 photoisomerisations/s (R*/s). Moving stimuli were designed and generated using a custom stimulus design and presentation GUI (StimGen) in either a MATLAB environment with Psychtoolbox or in a python environment with PsychoPy (https://github.com/benmurphybaum/StimGen).

### Electrophysiology

DSGCs were identified using 2-photon imaging in mouse lines with fluorescent labeling or were identified by their extracellular spiking responses in wild type mice. Electrodes were pulled from borosilicate glass capillaries to a resistance 3-6 MΩ. For extracellular recordings, electrodes were filled with Ringer’s solution. For voltage clamp recordings, electrodes contained the following in mM: 112.5 CH_3_CsO_3_S, 7.75 CsCl, 1 MgSO_4_, 10 EGTA, 10 HEPES, 5 QX-314-bromide (Tocris). For Ca^2+^ imaging current clamp recordings, electrodes were filled with the following in mM: 115 K-Gluconate, 7.7 KCl, 10 HEPES, 1 MgCl_2_, 2 ATP-Na_2_, 1 GTP-Na, 5 phosphocreatine, 2 QX-314, 0.2 Oregon Green Bapta-1, and 0.05 Sulphorhodamine 101. For some recordings, QX-314 was omitted and 1 µM TTX was included in the bath solution instead. Signals were sampled at 10 kHz and filtered at 2 kHz in a MultiClamp 700B amplifier (Molecular Devices). Analysis was performed using custom routines in MATLAB (Mathworks) and Igor Pro (Wavemetrics). The following pharmacological agents were added directly to the superfusion solution: DL-AP4 (50 µM; Tocris Bioscience), D-AP5 (50 µM; Abcam Biochemicals), UBP-310 (10 µM; Abcam Biochemicals), TTX (0.5-1 µM; Abcam Biochemicals), CNQX (20 µM; Tocris Bioscience).

### Calcium Imaging

For Ca^2+^ imaging experiments, DSGCs were patch clamped in current clamp mode and filled with the calcium indicator Oregon Green Bapta-1 (0.2 mM; ThermoFisher Scientific). For some experiments, a Ca^2+^-insensitive red dye, Sulforhodamine 101 (50 µM; ThermoFisher Scientific), was also included to help visualize the dendritic arbor. After waiting 15-20 minutes for the indicator to fill the dendrites, the microscope was focused onto the ON-stratifying dendritic layer. 2-photon excitation was delivered using an Insight DeepSee^+^ laser (Spectra Physics) tuned to 920 nm, guided by XY galvanometer mirrors (Cambridge Technology). Image scans were acquired using custom software developed by Dr. Jamie Boyd (University of British Columbia) in the Igor Pro environment (http://svn.igorexchange.com/viewvc/packages/twoPhoton/).

The laser power was rapidly modulated using a Pockels cell (Conoptics), such that laser power was extinguished during mirror flyback phases but was turned on during data acquisition phases. A counterphase signal was used to turn on and off the projector LEDs during flyback and scan phases, respectively. This method prevents the bright projector light from contaminating the fluorescence signal from the Ca^2+^ indicator, and also minimizes the time that the tissue is exposed to the laser.

PMT single photon currents were converted to voltages with a decay time constant of approximately 250 microseconds. These voltages were digitally sampled at a rate of 40 MHz, integrated for 1 microsecond through a custom time-integrating circuit (designed by Mike Delsey and Kerry Delaney, University of Victoria). This circuit outputs an analog voltage which was then digitized at 1 MHz (PCI-6110, National Instruments) for image formation. This helped decrease fluctuations in signal amplitude that arise from the asynchronous arrival times of single photons during the 1 microsecond dwell time at each image pixel, thereby improving signal to noise at low light levels characteristic of sparse synaptic activity.

### SAC ablation experiments

ON SACs on the null-side of the DSGCs, within 50-150 µm of its soma, were identified and targeted for physical ablation in ChaT-cre:nGFP mouse line. ~6-8 identified SACs were mechanically ablated by injecting 20 nA current for 10-15 s until the SAC cell membrane was ruptured (Jacoby et al., 2015). Ca^2+^ signals in the ON arbor of the targeted ON-OFF DSGC were then measured in response to preferred and null motion. DSIs were calculated according to Equation 6 (see *Quantification and Statistical Analysis*). To compare these values to previous control experiments, control DSIs were recalculated using only preferred and null responses, rather than using all 8 recorded directions. In some experiments, Ca^2+^ responses were measured in DSGC dendritic arbors before and after SAC ablation. In these experiments, DSGCs were loaded with OGB via single cell electroporation (Ding et al., 2016), and responses were measured in the presence of tetrodotoxin (0.5 µM TTX).

### Computational Modeling

A multi-compartmental model was coded in the NEURON environment(Hines and Carnevale, 1997). 177 pairs of excitatory and inhibitory synaptic inputs were distributed across the dendritic arbor of a reconstructed ON-OFF DSGC (Poleg-Polsky and Diamond, 2016). Membrane capacitance and axial resistance were set to 1 µF/cm^2^ and 100 Ω·cm respectively. Membrane channels and noise were modeled using a stochastic Hodgkin and Huxley distributed mechanism. Non-voltage-gated leak conductances were set to reverse at −60 mV. Active membrane conductances were placed at the soma, primary dendrites and terminal dendrites with the following densities in mS/cm^2^ (soma/primary dendrites/terminal dendrites): sodium (150/200/30), potassium rectifier (35/35/25), and delayed rectifier (0.8/0.8/0.8). This enabled the model DSGC to process inputs actively via dendritic sodium spikes. Sodium and potassium conductances were blocked for voltage-clamp recordings.

The membrane potential of 700 compartments across the dendritic arbor were recorded in response to a simulated edge moving across the receptive field at 1 mm/s. Onset times of each synaptic site varied around a mean value determined by the time at which the simulated edge passed over it. Random selections from Gaussian distributions determined whether or not synaptic release of inhibition and/or excitation was successful. For inhibition, the limits defining successful or unsuccessful release depend on the direction of motion, with more stringent limits during preferred motion when the probability of release is low, and broader limits during null motion when the probability of release is high. The release probability for excitatory synapses was held constant across all directions.

### Quantification and Statistical Analysis

All analysis and statistical comparisons were done using Igor Pro (WaveMetrics). All population data was expressed as mean ± standard deviation unless otherwise specified. Angular data was expressed as angular mean ± angular standard deviation (σ_θ_). The preferred direction and strength of the directional tuning was defined as the vector summation of the component responses for the 8 stimulus directions, which was computed as:

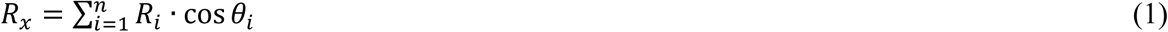

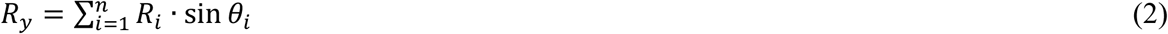

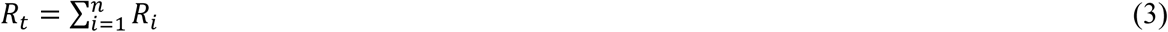

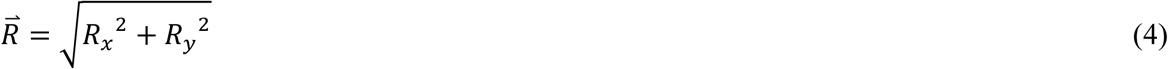

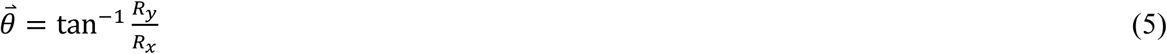

where *R*_*i*_ is the response for the *i*^th^ stimulus direction (*θ*), *R*_*x*_ and *R*_*y*_ are the *x* and *y* components of the response, and *R*_*t*_ is the summed total of all responses. 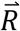 and 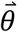 are the length of the vector sum and its angle, respectively. The direction selective index (DSI) is computed as the normalized length of the vector sum:

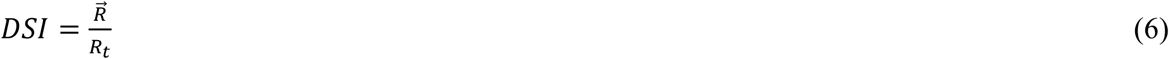

The angular standard deviation of a population of angles was calculated as:

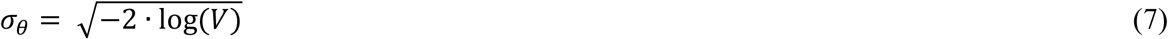

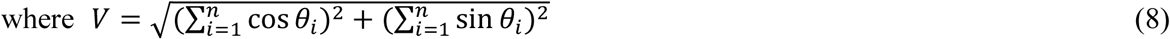

and *θ* is the *i*^th^ angle in the population.

Statistical comparisons between mean angles were done using the Watson-Williams test, and comparisons between the angular variance of two populations were done using the Angular Distance test, which is a Mann-Whitney-Wilcoxon test on the angular distances of each sample from its mean angle. Comparisons between mean DSIs over trials were done using the Student’s t test, while comparisons of distributions of DSIs (e.g. among many dendritic sites) were done using the Kolmogorov-Smirnov test. All other paired comparisons were done using the Student’s t test. All comparisons were done using two-tailed statistical tests. Differences were considered significant for p < 0.05.

## Data and Software Availability

The data analysis code is available upon request from the Lead Contact. The visual stimulation software (StimGen) is freely available online (https://github.com/benmurphybaum/StimGen).

## Contact for reagent and resource sharing

Further information and requests for resources and reagents should be directed to, and will be fulfilled by, the Lead Contact, Gautam B. Awatramani.

---

**Figure 6. Supplemental Figure 1.**
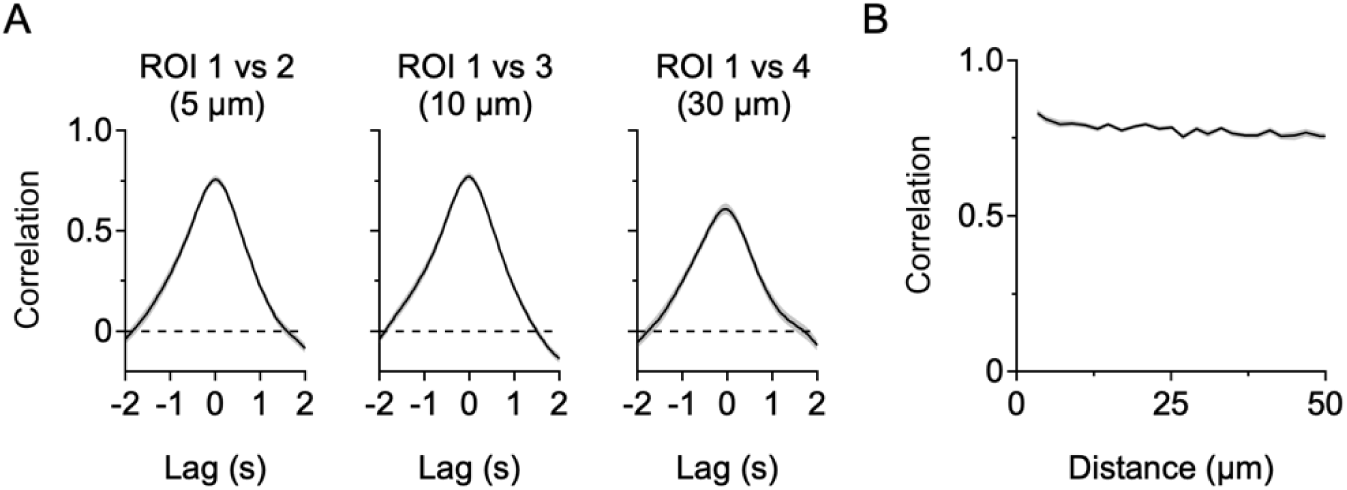
Stimulus correlations between DSGC dendritic sites. **A**, Correlations between the raw Ca^2+^ signals from the ROIs shown in **Figure 6** (without mean subtraction). **B**, Correlation between ROIs as a function of cable distance. Here, since Ca^2+^ signals arise at all sites during the period of stimulus presentation, the spatial decay in stimulus (signal) correlations is minimal.

**Figure 6. Supplemental Figure 2.**
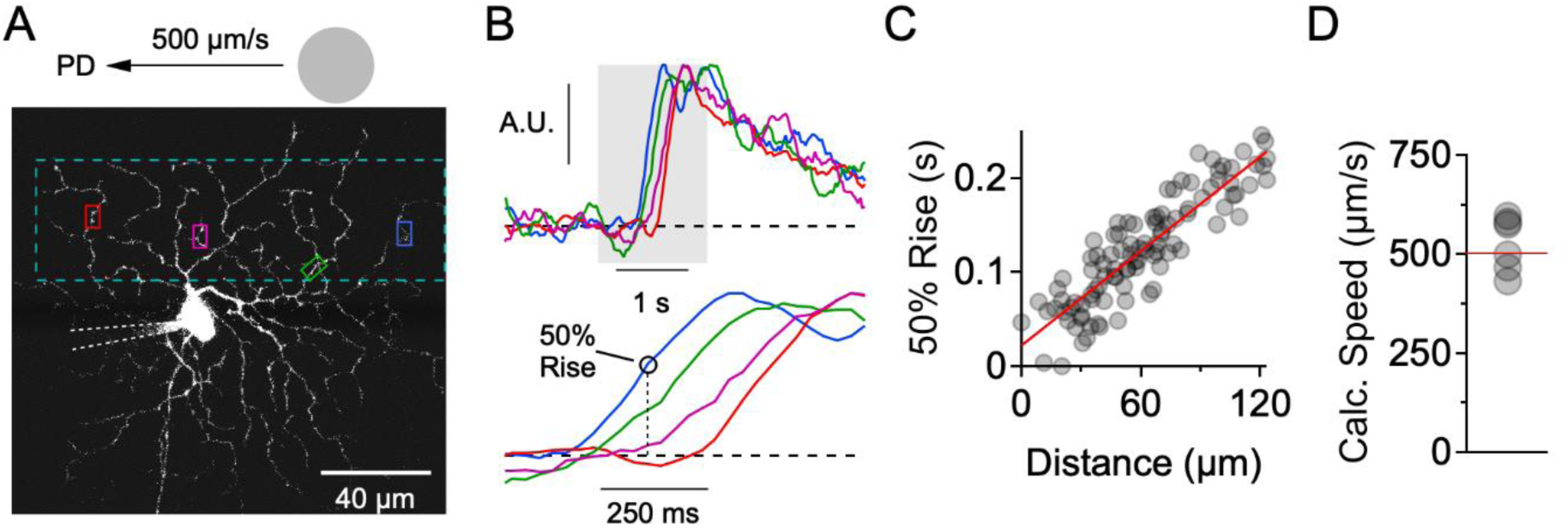
Preferred motion sequentially activates dendrites in the presence of NaV blockers. **A**, Image of the ON-stratifying dendritic arbor of an ON-OFF DSGC, which was filled with the Ca^2+^ indicator Oregon Green Bapta-1 (200 µM) through a patch clamp electrode. The NaV channel blocker QX-314 (2 mM) was also included in the electrode. **B**, Ca^2+^ signals for the four ROIs demarcated in **A**. Top, Signals were normalized so as to examine their relative timing onsets. Bottom, Expanded view of the shaded region, showing clear differences in onset time at the different ROIs. **C**, 50% rise time versus lateral distance along the stimulus trajectory for many small ROIs that were placed throughout the dendritic tree. The red line is a linear fit to the data. **D**, The slope of the linear fit predicts the speed of the stimulus (524 ± 28 µm/s), which was set to be 500 µm/s. This data set included DSGCs that were recorded with (n = 3) and without (n = 3) the presence of the NMDA receptor antagonist D-AP5 (50 µM). No differences were found in terms of the onset timing of the Ca^2+^ signals, so the data sets were combined.

**Figure 6, Supplemental Figure 3.**
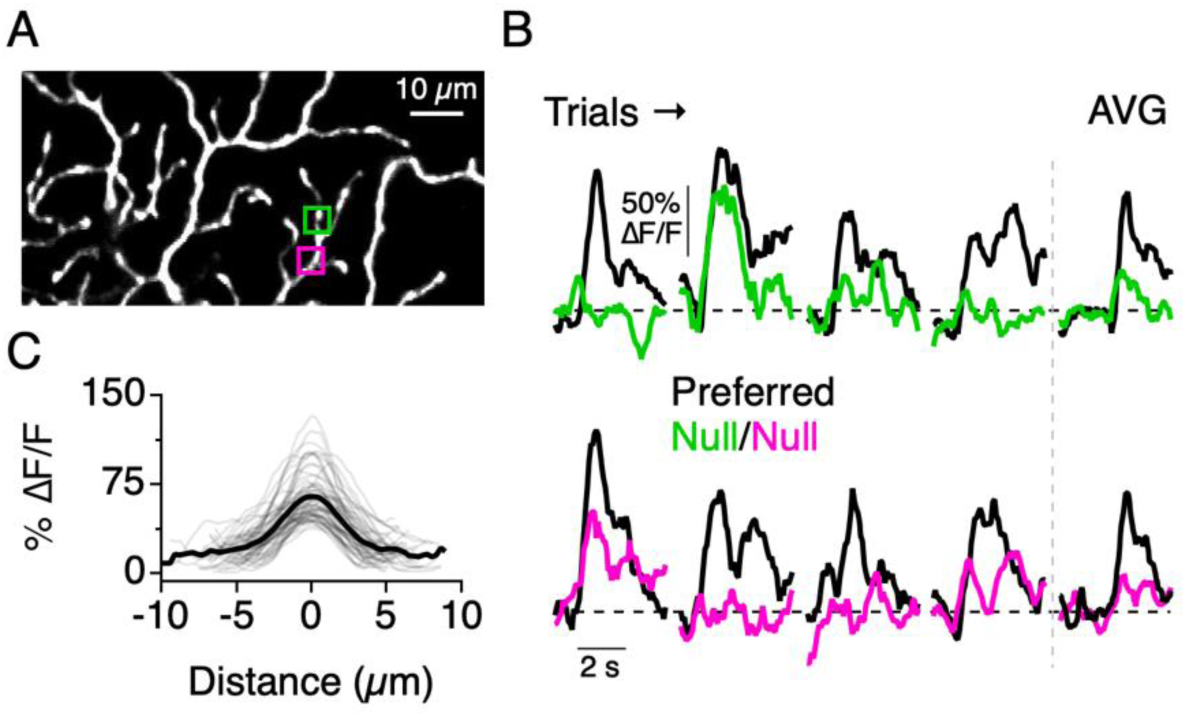
Sparse Ca^2+^ signals in response to null motion. **A**, Imaging area in the ON arbor of an ON-OFF DSGC, and two small ROIs used for analysis. **B**, Ca^2+^ signals from the ROIs demarcated in **A**, in response to preferred and null motion over multiple trials. Right, Average signals over four trials. Ca^2+^ signals were largely suppressed in response to null motion, but occasionally dendritic sites would show activity. **C**, Line profiles of the ΔF/F across active dendritic sites. Activity in response to null motion was generally limited to small dendritic regions. Solid line is the average line profile, and the thin lines are from individual ROIs. Gaussian fits to the line profiles had an average width of 3.0 ± 1.2 µm (µ ± s.d., n = 61 ROIs from 7 cells).

